# Uncoupling splicing from transcription using antisense oligonucleotides reveals a dual role for I exon donor splice sites in antibody class switching

**DOI:** 10.1101/850867

**Authors:** Anne Marchalot, Mohamad Omar Ashi, Jean-Marie Lambert, Nivine Srour, Laurent Delpy, Soazig Le Pennec

## Abstract

Class switch recombination (CSR) changes antibody isotype by replacing Cμ constant exons with different constant exons located downstream on the immunoglobulin heavy (*IgH*) locus. During CSR, transcription through specific switch (S) regions and processing of noncoding germline transcripts (GLTs) are essential for the targeting of Activation-Induced cytidine Deaminase (AID). While CSR to IgG1 is abolished in mice lacking Iγ1 exon donor splice site (dss), many questions remain regarding the importance of I exon dss recognition in CSR. To further clarify the role of I exon dss in CSR, we first evaluated RNA polymerase II (RNA pol II) loading and chromatin accessibility in S regions after activation of mouse B cells lacking Iγ1 dss. We found that deletion of Iγ1 dss markedly reduced RNA pol II pausing and active chromatin marks in the Sγ1 region. We then challenged the post-transcriptional function of I exon dss in CSR by using antisense oligonucleotides (ASO) masking I exon dss on GLTs. Treatment of stimulated B cells with an ASO targeting Iγ1 dss, in the acceptor Sγ1 region, or Iμ dss, in the donor Sμ region, did not decrease germline transcription but strongly inhibited constitutive splicing and CSR to IgG1. Altogether, this study reveals that the recognition of I exon dss first supports RNA pol II pausing and the opening of chromatin in targeted S regions and that GLTs splicing events using constitutive I exon dss appear mandatory for the later steps of CSR, most likely by guiding AID to S regions.

## INTRODUCTION

During immune responses, B cells can diversify the immunoglobulin (Ig) repertoire through class switch recombination (CSR) and somatic hypermutation (SHM). SHM introduces mutations in the variable (V) regions of Ig genes modifying antibody affinity for a cognate antigen. The mouse Ig heavy chain (*IgH*) locus is comprised of eight constant genes (C_H_). CSR involves long-range interactions at the *IgH* locus and occurs between GC-rich repetitive switch (S) DNA regions preceding each C_H_ gene due to the enzymatic activity of Activation-Induced Deaminase (AID) ^1,2^. Thus, CSR replaces the Cμ exons by a downstream constant gene Cγ, Cε or Cα allowing expression of antibodies with different isotypes (from IgM to IgG, IgE or IgA) and effector functions.

Transcription through S regions is required for CSR ^3^. Germline (GL) transcription is initiated from intervening (I) promoters a few kilobases upstream of both the donor and the acceptor S regions. The μ I promoter drives constitutive transcription through Sμ while γ, ε or α I promoters are inducible. Primary GL transcripts (GLTs) exhibit a conserved structure composed of a noncoding I exon, an intronic S region and C_H_ exons ^4,5^. In mature GLTs, the I exon is spliced to the first exon (CH1) of the adjacent constant gene. Because multiple stop codons are present in the three reading frames of I exons, these GLTs do not encode peptides of significant lengths.

During CSR, AID initiates double-strand DNA breaks (DSBs) by deaminating cytidines inside the transcribed S regions. GL transcription through S regions of C_H_ gene favors AID accessibility to S regions ^3^. GL transcription promotes generation of RNA:DNA hybrid structures (R-loops) ^6,7^ revealing single-stranded DNA (ssDNA) that serves as substrate for AID ^8^. The impairment of transcription elongation upon R-loop formation ^9^ may favor RNA polymerase II (RNA pol II) pausing. RNA pol II pausing then promotes AID recruitment to S regions ^10,11^. Paused RNA pol II and histone modifications associated with “open” chromatin, such as histone H3 lysine 4 trimethylation (H3K4me3) and histone H3 lysine 9 acetylation (H3K9ac), are enriched in transcribed I-S regions and have been involved in AID targeting to S regions primed for CSR ^12–16^. Moreover, the Suppressor of Ty 5 homolog (Spt5) transcription elongation factor and the RNA exosome, a cellular RNA-processing degradation complex, associate with AID together with paused RNA pol II in transcribed S regions and are required for CSR ^17,18^.

Beyond the prerequisite transcription of S regions, splicing of GLTs has been proposed to be important for the CSR process. Notably, CSR to IgG1 is severely impaired in a mouse model lacking the Iγ1 exon donor splice site (dss) ^19,20^. Further supporting a role for splicing of GLTs in CSR, several RNA processing and splicing factors are critical regulators of CSR ^21,22^. Interestingly, it has recently been proposed that intronic switch RNAs produced by the splicing of primary GLTs act as guide RNAs and target AID to DNA in a sequence-specific manner ^23^. After lariat debranching by the RNA debranching enzyme (DBR1), these switch RNAs are folded into G-quadruplexes. G-quadruplexes and AID are targeted to S region DNA through post-transcriptional action of the DEAD-box RNA helicase 1 (DDX1) ^24^.

Even though these data suggest that processing of GLTs by the splicing machinery is necessary for CSR, the precise role of I exon dss recognition in antibody class switching remains largely unknown. To address this issue, we first analysed whether the presence of Iγ1 exon dss could influence RNA pol II pausing and chromatin accessibility of Sγ1 region, as early events leading to CSR to IgG1. For that, Chromatin Immunoprecipitation (ChIP) experiments were performed in stimulated B cells from the previously described human MetalloThionein II_A_ (*hMT*) and *s-hMT* (splice hMT) mouse models, lacking or harbouring Iγ1 exon dss, respectively ^19,20^. We next specifically evaluated the impact of GLTs splicing on CSR to IgG1 by using antisense oligonucleotides (ASOs) targeting specific I exon dss on primary GLTs, from both donor and acceptor S regions. Contrary to the models used previously to study the impact of I exons on CSR, treatment of mouse B cells by ASOs masks only a short RNA sequence (23 to 25 nucleotides) surrounding the I exon dss on primary GLTs. This antisense strategy bypassing the impact of I exon dss recognition on transcription is very useful for studying the involvement of I exon dss recognition in CSR at the post-transcriptional level. Collectively, our data indicate that the recognition of I exon dss exerts both transcriptional and post-transcriptional roles during CSR.

## RESULTS

### RNA pol II pausing and histone modifications upstream of the Sγ1 region are altered in mice lacking Iγ1 dss

To clarify the role of I exon dss in CSR, we first performed comparative experiments in homozygous *s-hMT* and *hMT* mice ^19,20^. In these models, the promoter and Iγ1 exon have been replaced by the lipopolysaccharide (LPS)-inducible *hMT* promoter and an artificial I_hMT_ exon containing (*s-hMT*) or lacking (*hMT*) Iγ1 dss (Figure 1A). As previously described by Lorenz and collaborators ^20^, IgG1 class switching was almost abolished in *hMT* mice, whereas CSR to other Ig isotypes remained unaffected (Supplementary figure 1A-C). This was not associated with a difference in AID mRNA expression between splenic B cells isolated from *hMT* and *s-hMT* mice (Supplementary figure 1D). Previous nuclear run-on assays have shown that the transcription rate of the Sγ1 region was higher in LPS-stimulated B cells from *s-hMT* mice than from *hMT* mice ^20^, suggesting a role for Iγ1 exon dss in GL transcription of the Sγ1 region. In agreement with these results, we found low levels of unspliced γ1 GLTs in LPS-stimulated B cells from *hMT* mice, compared to *s-hMT* (Figure 1B).

**Figure 1.**
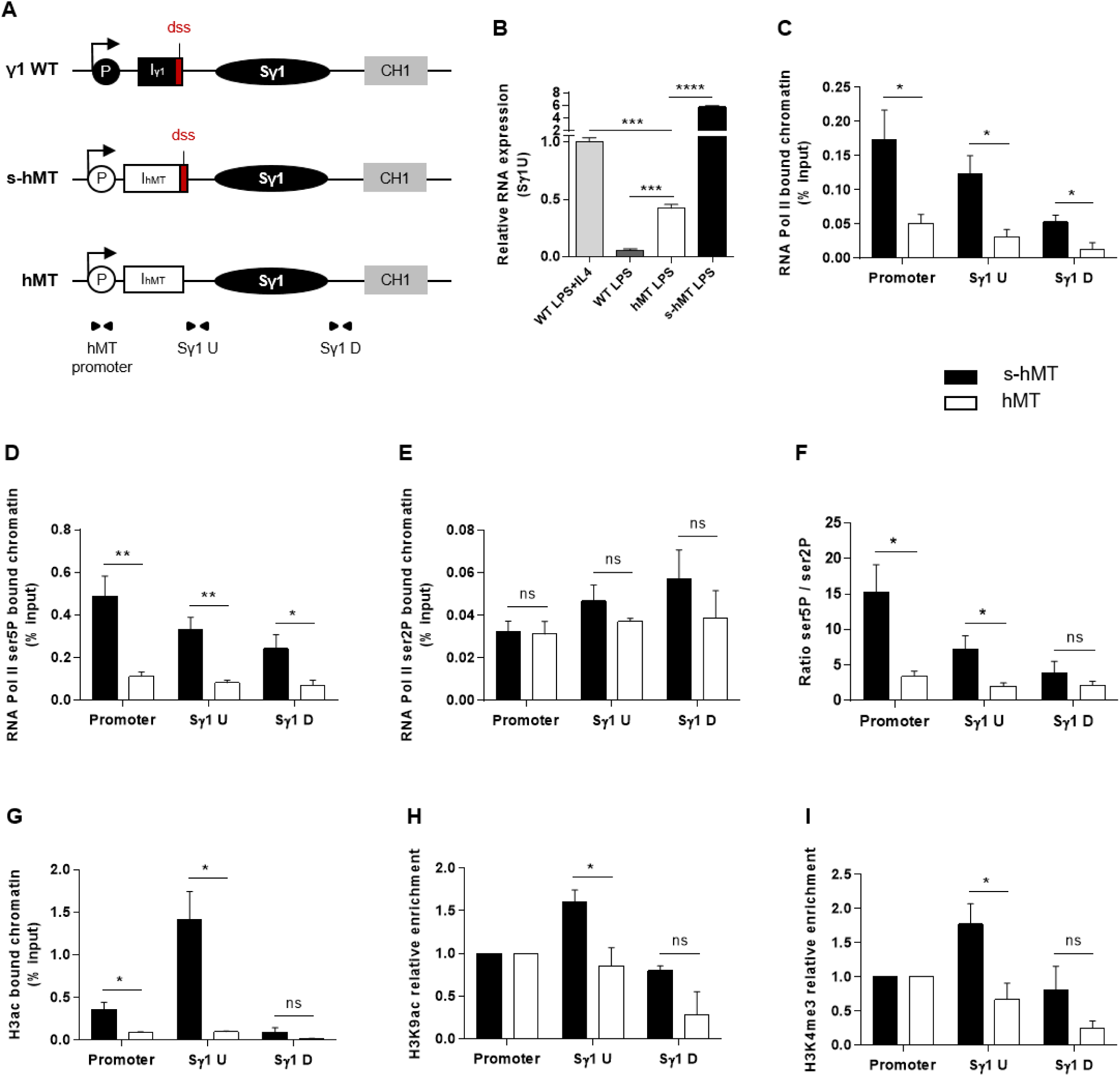
Deletion of Iγ1 exon dss decreased RNA pol II pausing and chromatin accessibility in the Sγ1 region. (A) Wild type (WT) γ1 locus and *s-hMT* and *hMT* murine models. Black and grey elements represent gene segments of the γ1 WT locus. The white elements are gene segments specific to *s-hMT* and *hMT* models and are not present in WT animals. The red fragment corresponds to the sequence containing the I exon donor splice site (dss), retained in the *s-hMT* model and absent in the *hMT* model. P, promoter; Iγ1, γ1 I exon; I_hMT_, hMT I exon; Sγ1, γ1 switch region; CH1, γ1 constant exon 1. (B) Splenic B cells were isolated from WT, homozygous *s-hMT* and homozygous *hMT* mice and stimulated with LPS + IL4 or LPS. After 2 days stimulation, γ1 GLTs expression relative to GAPDH mRNA expression was monitored by quantitative RT-PCR using Sγ1U primers described in schema A. Expression of γ1 GLTs in B cells isolated from WT mice and stimulated by LPS + IL4 was normalized to 1. (C-I) Splenic B cells were isolated from homozygous *s-hMT* and *hMT* mice and stimulated with LPS. After 2 days stimulation, the cells were analysed for RNA pol II (C), Ser5P RNA pol II (D) and Ser2P RNA pol II (E) levels in the γ1 locus by ChIP coupled to quantitative PCR. Ser5P RNA pol II / Ser2P RNA pol II ratios are indicated (F). Cells were also analysed for H3ac (G), H3K9ac (H) and H3K4me3 (I) levels in the γ1 locus by ChIP coupled to quantitative PCR. Background signals from mock samples with irrelevant antibody were subtracted. Values were normalized to total input DNA. Relative variations of H3K9ac and H3K4me3 marks were expressed after normalization to the value obtained for the promoter. Position and sequence of primers used for quantitative PCR is described in schema A (triangles) and Supplementary table 1. (B-I) Data are means ± SEM of at least two independent experiments, n=3 to 5 for each genotype. Unpaired two-tailed Student’s t test was used to determine significance. ns: non significant; *, P < 0.05, **, P < 0.01, *** P <0.001, **** P <0.0001.

Paused RNA pol II and histone modifications associated with “open” chromatin are involved in AID recruitment to S regions during CSR ^10,12,13,15,16,18^. To study the role of I exon dss in RNA pol II pausing and chromatin remodelling in the Sγ1 region, we performed ChIP experiments in splenic B cells isolated from *hMT* and *s-hMT* mice after 2 days LPS stimulation. First, antibodies directed against total RNA pol II, elongating RNA pol II (Ser2P) or pausing RNA pol II (Ser5P) were used. In agreement with the low level of γ1 GLTs described in Figure 1B, stimulated B cells from *hMT* mice displayed significantly decreased total RNA pol II loading throughout the Sγ1 region, compared to *s-hMT* mice (Figure 1C). Interestingly, a significant decrease in Ser5P RNA pol II loading (Figure 1D), but not Ser2P RNA pol II loading (Figure 1E), was observed in the promoter-Sγ1 region in stimulated B cells from *hMT* compared to *s-hMT* mice. Accordingly, stimulated B cells from *s-hMT* mice exhibited significantly higher Ser5P/Ser2P ratios upstream from the Sγ1 region than B cells from *hMT* animals (Figure 1F), suggesting that the recognition of I exon dss regulated RNA pol II pausing upstream from the Sγ1 region. As a control, we found similar Ser5P and Ser2P RNA pol II loading in Sμ and Sγ2b regions in stimulated B cells from both *hMT* and *s-hMT* mice (Supplementary figure 2).

We next performed similar ChIP experiments using antibodies directed against variants of histone H3. We observed very low levels of acetylated histone H3 (H3ac), a general marker of chromatin accessibility, throughout the whole promoter-Sγ1 region in LPS-stimulated B cells from *hMT* compared to *s-hMT* mice (Figure 1G). In addition, it has been previously shown that H3K9ac and H3K4me3 levels in the region upstream from Sγ1 were very low in LPS-stimulated B cells from *hMT* mice, compared to LPS + interleukin 4 (IL4)-stimulated B cells from wild type (WT) mice ^25^. Therefore, to evaluate the specific enrichment of active histone marks in the Sγ1 region in *hMT* and *s-hMT* models, H3K9ac and H3K4me3 levels were expressed as fold change after normalisation to values obtained at the promoter. In B cells from *s-hMT* mice, the highest levels of H3ac, H3K9ac and H3K4me3 histone forms were observed in the region upstream from Sγ1 (Figure 1G-I). In contrast, this increase was not anymore observed in B cells from *hMT* mice.

Collectively, our data indicate that Iγ1 exon dss recognition is of key importance for transcriptional activity of the Sγ1 region by supporting RNA pol II pausing and chromatin remodelling, two occurrences required for AID recruitment in S regions and efficient CSR.

### ASO strategy targeting I exon donor splice site allows uncoupling germline transcription and splicing in wild type B cells

We next wanted to study the function of I exon splicing on CSR. For this purpose, we treated stimulated splenic B cells isolated from WT mice with vivo-morpholino antisense oligonucleotide targeting the dss sequence of Iγ1 exon (Iγ1 dss ASO) (Figure 2A). Indeed, we previously demonstrated that passive administration of these Phosphorodiamidate Morpholino Oligonucleotides complexed with an octaguanidine dendrimer could very efficiently modulate the splicing of Ig transcripts ^26^. First, we wanted to verify that masking the dss sequence of Iγ1 exon on GLTs by ASO did not inhibit γ1 GL transcription. After 2 days culture with LPS + IL4 and 2 μM ASO, expression of unspliced γ1 GLTs in activated B cells was significantly increased after treatment with Iγ1 dss ASO (Figure 2B), compared to cells treated with an irrelevant control ASO. This suggested an accumulation of unspliced γ1 GLTs due to Iγ1 dss ASO-induced splicing inhibition. In order to study the consequences of Iγ1 dss ASO on spliced γ1 GLTs, we next performed PCR using the Iγ1-Cγ1 primer pair (Iγ1-for and Cγ1-rev, described in Figure 2C and Supplementary table 1) on cDNA from ASO-treated WT splenic B cells stimulated by LPS + IL4. After 2 days, the γ1 GLTs profile was strikingly different in control and Iγ1 dss ASO conditions (Figure 2C-up). A band corresponding to the constitutively spliced γ1 GLT (involving the constitutive Iγ1 exon dss) was strongly detected in control but slightly detected upon Iγ1 dss ASO treatment (Figure 2C-up and Supplementary figure 3). Remarkably, in both control and Iγ1 dss ASO conditions, additional bands were also detected that could account for new splicing isoforms. Sanger sequencing indeed revealed alternative transcripts so far not described in the literature (alternative transcripts 1, 2 and 3; Figure 2C-middle and Supplementary figure 3) involving donor and acceptor splice sites internal to the Iγ1 exon. The fact that these alternative splice sites were predicted with high consensus value by the HSF 3.1 tool (HSF 3.1, 24/07/2019, http://www.umd.be/HSF/HSF.shtml, ^27^) (Figure 2C-down) indicated that the different PCR products were *bona fide* spliced transcripts. As a proof of ASO efficiency, treatment with Iγ1 dss ASO almost abolished the major constitutive GLT and prevented alternative transcript 1 detection (Figure 2C-up). Thus, after treatment with Iγ1 dss ASO, transcript isoforms using the constitutive Iγ1 dss represented only 20% of the detected transcripts (*versus* 90% in control). Another consequence of Iγ1 dss ASO treatment is the detection of alternative transcript 3 and a better detection of alternative transcript 2.

**Figure 2.**
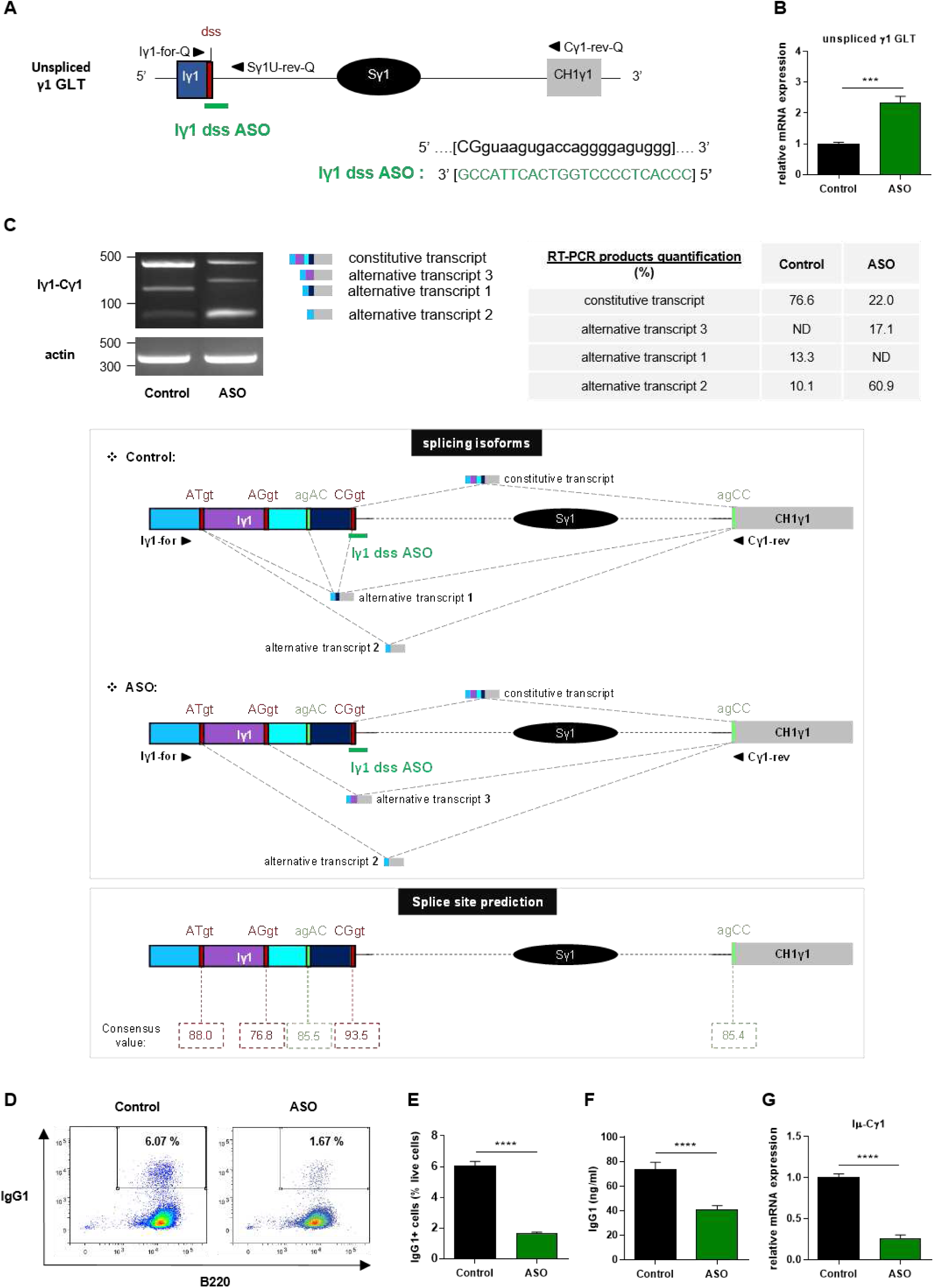
Treatment with Iγ1 exon dss ASO inhibited γ1 GLTs constitutive splicing and IgG1 class switching in B cells. (A) Antisense oligonucleotide targeting the donor splice site of Iγ1 exon (Iγ1 dss ASO) was designed and synthetized as “vivo-morpholino ASO” permitting passive administration of ASO in cells (Gene Tools, LLC). Targeted γ1 GLT (uppercase: exon sequence; lowercase: intron sequence) and Iγ1 dss ASO sequences are indicated. Iγ1, γ1 I exon; Sγ1, γ1 switch region; CH1γ1, γ1 constant exon 1; dss, donor splice site. (B-G) Splenic B cells were isolated from *C57BL/6* mice, stimulated with LPS + IL4 and treated with 2 μM Iγ1 dss ASO (ASO) or an irrelevant ASO (control) during 2 days (B-C) or 3 days (D-G). (B) Unspliced γ1 GLTs expression relative to GAPDH mRNA expression was monitored by quantitative RT-PCR using Iγ1-for-Q and Sγ1U-rev-Q primers described in schema A. Expression of γ1 GLTs in control B cells was normalized to 1. (C-up) RT-PCR was performed using Iγ1-for and Cγ1-rev primers (position described in schema C-middle) to identify constitutively and alternatively spliced transcripts. PCR products were analysed on agarose gels. Expression of actin mRNA is also shown. Molecular markers in base pairs are indicated. Schematic representation of the different γ1 spliced transcripts is indicated on the right and transcript sequences are given in Supplementary figure 3. One experiment out of three is shown. Quantification of amplification products was done using an Agilent Bioanalyzer. Data are expressed as percentage of each isoforms among the detected transcripts. ND: not detected. (C-middle) Schematic representation of γ1 spliced transcripts detected in B cells from *C57BL/6* mice after treatment with an irrelevant ASO (control) or Iγ1 dss ASO (ASO). Grey hatched lines represent splicing events involving constitutive and alternative splice sites. Donor and acceptor splice sites are indicated in red and green respectively. (C-down) Consensus value (ranging from 0 to 100) of each predicted splice site determined using HSF 3.1 tool. (D) Flow cytometry analysis of purified B cell populations using the indicated cell surface markers. Plots are gated on live cells. The percentage of B220^+^IgG1^+^ cells is indicated. One experiment out of five is shown. (E) Percentage of IgG1 positive cells determined by flow cytometry. (F) Quantification of IgG1 in culture supernatants by ELISA. (G) Post-switch Iμ-Cγ1 mRNA expression relative to GAPDH expression was monitored by quantitative RT-PCR. Expression of post-switch mRNAs in control B cells was normalized to 1. Sequence of primers used for RT-PCR and quantitative RT-PCR are indicated in Supplementary table 1. (B and E-G) Data are means ± SEM of two independent experiments, n=5 to 8 for each group. Unpaired two-tailed Student’s t test was used to determine significance. *** P <0.001, **** P <0.0001.

Since our data showed that Iγ1 dss ASO strongly decreased γ1 GLT constitutive splicing, ASO strategy is a good tool to study the function of Iγ1 exon splicing on CSR.

### Treatment with Iγ1 exon dss ASO specifically inhibits IgG1 class switching

We next investigated the effect of ASO-mediated γ1 GLTs splicing inhibition on CSR to IgG1. After 3 days culture with LPS + IL4 and 2 μM ASO, CSR to IgG1 was greatly diminished in B cells treated with Iγ1 dss ASO compared to B cells treated with an irrelevant control ASO. The percentage of IgG1 positive B cells determined by flow cytometry was decreased 4-fold in Iγ1 dss ASO-treated B cells (Figure 2D-E) and IgG1 levels determined in culture supernatants by Enzyme Linked Immuno Sorbent Assay (ELISA) were almost 2-fold lower in Iγ1 dss ASO-treated cells (Figure 2F) than in control B cells. In agreement with the CSR defect, Iμ-Cγ1 switched transcripts were decreased by 4 fold in Iγ1 dss ASO-treated B cells (Figure 2G). In order to control Iγ1 dss ASO specificity, we realized similar experiments in B cells stimulated by anti-CD40 + IL4. Similarly to what observed in LPS + IL4 -stimulated B cells, after 2 days culture with anti-CD40 + IL4 and 2 μM ASO, expression of unspliced γ1 GLTs was significantly increased (Figure 3B) and CSR to IgG1 was significantly diminished (Figure 3C-D) in B cells treated with Iγ1 dss ASO compared to B cells treated with an irrelevant control ASO. However, expression of unspliced Iε GLTs (Figure 3E) and CSR to IgE (Figure 3F-G) were similar in B cells treated with the irrelevant control ASO or Iγ1 dss ASO. This showed that Iγ1 dss ASO specifically decreased CSR to IgG1 by inhibiting splicing precisely on γ1 GLTs.

**Figure 3.**
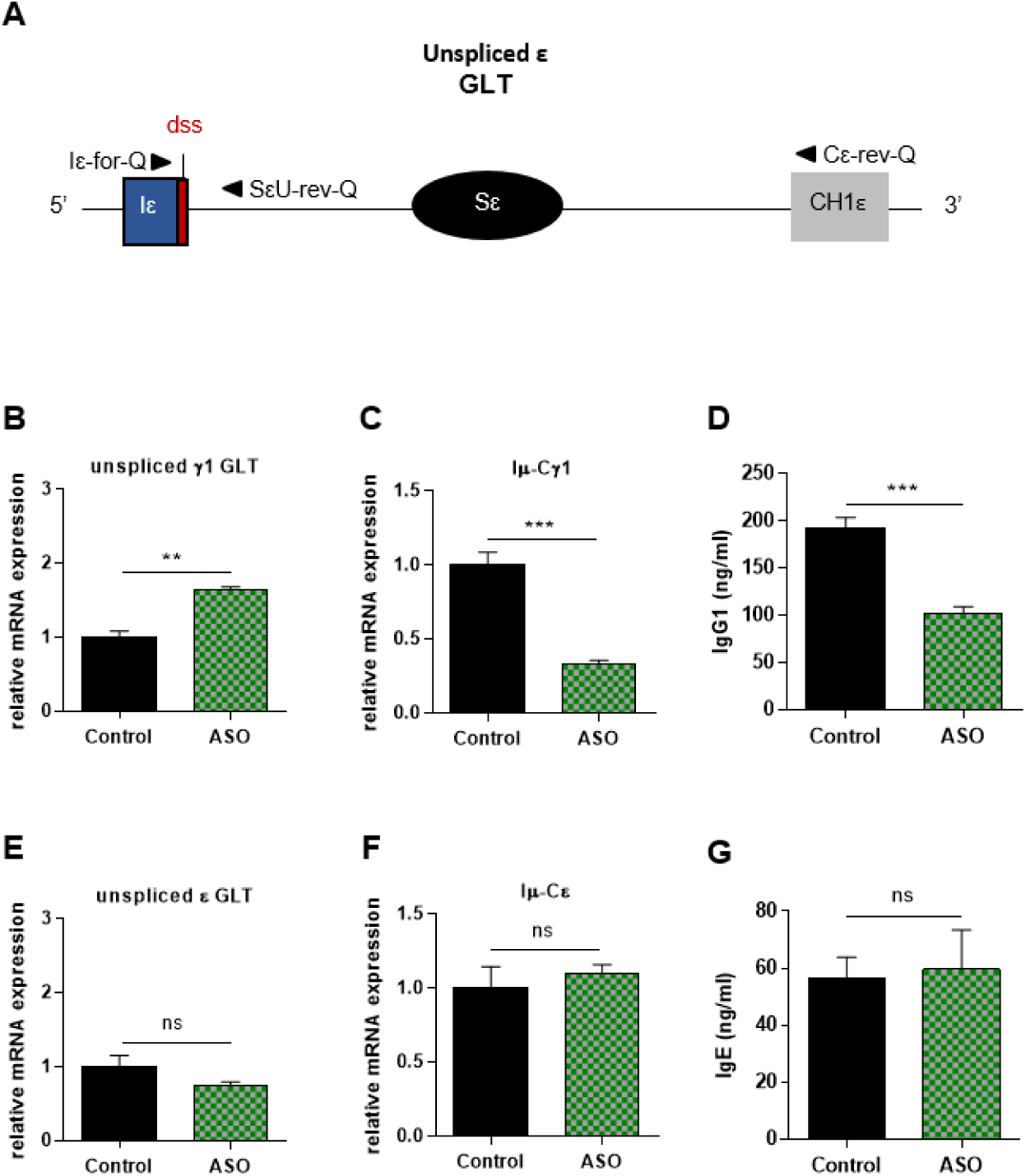
Specific IgG1 class switching inhibition in B cells treated by Iγ1 exon dss ASO. Splenic B cells were isolated from *C57BL/6* mice, stimulated with anti-CD40 + IL4 and treated with 2 μM Iγ1 dss ASO (ASO) or an irrelevant ASO (control) during 2 days (B, E) or 4 days (C, D, F, G). (A) Schematic representation of unspliced ε GLT. Iε, ε I exon; Sε, ε switch region; CH1ε, ε constant exon 1; dss, donor splice site. (B, E) Unspliced γ1 (B) and ε (E) GLTs expression monitored as described in figure 2. Iε-for-Q and SεU-rev-Q primers, described in schema A, were used for ε GLTs expression determination. (C, F) Post-switch Iμ-Cγ1 and Iμ-Cε mRNA expression monitored as described in figure 2. (D, G) Quantification of IgG1 and IgE in culture supernatants by ELISA. (B-G) Data are means ± SEM of two independent experiments, n=3 to 4 for each group. Unpaired two-tailed Student’s t test was used to determine significance. ns: non significant, ** P < 0.01, *** P <0.001.

Collectively, these data indicate that splicing of γ1 GLTs is necessary for CSR to IgG1 and strongly suggest that the use of the constitutive Iγ1 exon dss during splicing is required for efficient CSR.

### Treatment with an ASO targeting the constitutive I exon dss in the donor Sμ region also inhibits IgG1 class switching

Our results showed that ASO strategy targeting the constitutive I exon dss from Sγ1 acceptor region inhibited CSR to IgG1. We next developed a similar approach to determine whether specifically masking the constitutive I exon dss from the donor Sμ region could also inhibited CSR to IgG1. Splenic B cells isolated from WT mice were stimulated by LPS + IL4 and treated with an ASO targeting the constitutive Iμ exon dss (Iμ dss ASO) (Figure 4A). In order to study the consequences of Iμ dss ASO on spliced Iμ GLTs, we performed PCR using the Iμ-Cμ primer pair (Iμ-for and Cμ-rev, described in Figure 4A and Supplementary table 1) on cDNA from stimulated splenic B cells treated with 2 μM ASO for 2 days. A band corresponding to the constitutively spliced Iμ transcript (involving the constitutive Iμ exon dss) was detected in control but slightly detected after Iμ dss ASO treatment (Figure 4B). Similarly to what observed in stimulated splenic B cells treated with Iγ1 dss ASO, we detected the presence of alternatively spliced Iμ transcripts in cells treated with Iμ dss ASO. These alternative transcripts were indicative of the use of alternative Iμ dss present in the intronic S region previously described by Kuzin and collaborators ^28^. In addition, after 3 days culture with LPS + IL4 and 2 μM ASO, CSR to IgG1 was significantly diminished in B cells treated with Iμ dss ASO compared to B cells treated with an irrelevant control ASO (Figure 4C-D). After 2 days 2 μM ASO treatment, expression of unspliced γ1 GLTs was similar in B cells treated with an irrelevant control or Iμ dss ASO (Figure 4E). This showed that the defect in IgG1 class switching observed after treatment by Iμ dss ASO was not due to an inhibition of γ1 GLTs splicing but to the splicing decrease of Iμ GLTs.

**Figure 4.**
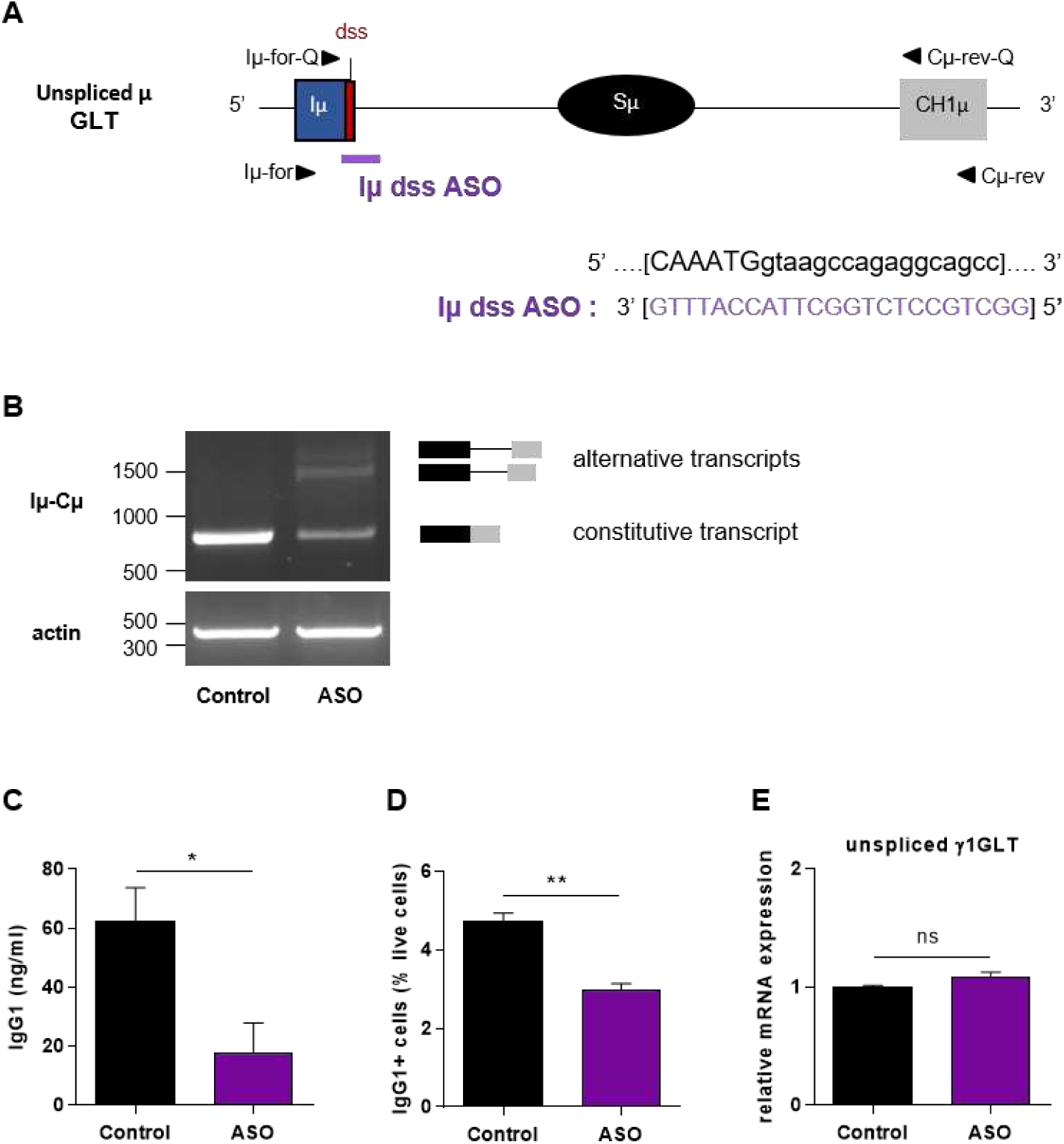
Defective IgG1 class switching in B cells treated by Iμ exon dss ASO. (A) Antisense oligonucleotide targeting the donor splice site of Iμ exon (Iμ dss ASO) was designed and synthetized as “vivo-morpholino ASO” permitting passive administration of ASO in the cells (Gene Tools, LLC). Targeted μ GLT (uppercase: exon sequence; lowercase: intron sequence) and Iμ dss ASO sequences are indicated. Iμ, μ I exon; Sμ, μ switch region; CH1μ, μ constant exon 1; dss, donor splice site. (B-E) Splenic B cells were isolated from *C57BL/6* mice, stimulated with LPS + IL4 and treated with 2 μM Iμ dss ASO or an irrelevant control ASO for 2 days (B, E) or 3 days (C, D). (B) Constitutively and alternatively spliced μ transcripts analysed as described in figure 2 using Iμ-for and Cμ-rev primers, described in schema A. (C) Quantification of IgG1 in culture supernatants by ELISA. (D) Percentage of IgG1 positive cells determined by flow cytometry as described in figure 2. (E) Unspliced γ1 GLTs expression monitored as described in figure 2. (C-E) Data are means ± SEM, n=3 for each group. Unpaired two-tailed Student’s t test was used to determine significance. ns: non significant, * P < 0.05, ** P <0.01.

Our results indicated that splicing of GLTs involving the constitutive Iμ dss is necessary for efficient CSR to IgG1. Therefore, correct splicing of GLTs produced in both the donor Sμ and the acceptor Sγ1 regions is required for efficient IgG1 class switching.

## DISCUSSION

S region transcription *per se* promotes basal class switch recombination but, for optimal efficiency, the process requires the presence of the intact I region, implicating factors beyond transcription through the S region in the regulation of class switching ^29^. Transcription and pre-mRNA processing are functionally coupled. In our study, through the experimental uncoupling of splicing from transcription, we identified distinct functions of I exon dss that control antibody class switching at transcriptional and post-transcriptional levels. During transcription, we provide evidence that deletion of the Iγ1 exon dss decreased accumulation of Ser5P RNA pol II and chromatin accessibility in the promoter-Sγ1 region. Moreover, our data demonstrated that, in S regions primed for CSR, GLTs splicing involving the constitutive I exon dss is an essential step for efficient CSR.

Regarding the impact of I exon dss on transcription, our comparative analysis of *s-hMT* and *hMT* B cells indicated that, in the absence of Iγ1 dss, GL transcription and RNA pol II loading were very weak at the γ1 locus. Interestingly, RNA pol II binding was strongly diminished at the *hMT* promoter, suggesting that the presence of Iγ1 exon dss enhanced the initiation of GL transcription. These data are consistent with the poor transcription observed upon deletion of a large portion of Iγ2b exon including dss ^30^. Although the function of I exon dss in transcription regulation has been overlooked, such intron-mediated enhancement of gene expression has long been described in a wide range of organisms, including mammals ^31^. Indeed, it has been demonstrated that the presence of a dss facilitates the transcription preinitiation complex assembly and stimulates transcription even in the absence of splicing ^32,33^. Whether the recognition of I exon dss promotes the formation of pre-initiation complex ^32^ and involves a gene looping interaction between promoter and dss, as described in yeast cells ^34^, remains to be investigated. Our ChIP analysis further indicated that, in the absence of Iγ1 exon dss, RNA pol II pausing is markedly decreased in the promoter-Sγ1 region, whereas the rate of RNA pol II elongation remains mostly unchanged. It is tempting to speculate that the I exon dss acts as an anchor for the co-transcriptional machinery and is required for appropriate associations of critical factors, like Spt5, with RNA pol II to enable recruitment of AID to S regions. After B cells stimulation, transcribed I-S regions are a focus for increased modified histones, such as H3ac, H3K9ac or H3K4me3, and chromatin accessibility ^13,16^. Reinforcing the idea that GL transcription, RNA pol II pausing and the establishment of active chromatin marks in S regions are interconnected ^10^, we found a global reduction of active H3ac marks in the promoter-Sγ1 region of stimulated B cells from *hMT* mice, compared to *s-hMT*. The H3K9ac and H3K4me3 enrichment specifically detected upstream the Sγ1 region was also lost in *hMT* mice. Interestingly, it has been proposed that H3K4me3 serves as a mark for recruiting the recombinase machinery for CSR independently of its function in transcription ^15,35^. These studies have shown that Spt5 and the histone chaperone FAcilitates Chromatin Transcription (FACT) are not required for transcription of S regions but regulate H3K4me3 modification and DNA cleavage in CSR ^15,35^.

Iγ1 exon dss recognition is necessary for the regulation of GL transcription. Consequently, using classical I exon deletion and/or replacement mouse models neither allow distinguishing the roles of I exon dss in the interconnected processes of transcription and splicing nor permit distinguishing between requirements of splicing *per se* from that of stable S GLTs ^36^. Several studies including ours have shown the efficacy of ASO-mediated approaches to modify RNA splicing in B-lineage cells ^26,37,38^. Here, we used ASOs targeting short sequence (23 to 25 nucleotides) spanning the Iγ1 or Iμ exon dss on pre-mRNA to analyse the consequences of GLTs splicing defect on CSR to IgG1 in splenic B cells from WT mice. We showed that, in addition to its positive effect on transcription, I exon dss recognition is required at the post-transcriptional level for efficient CSR to IgG1. Moreover, our results strongly suggest that splicing of GLTs *per se* is not sufficient to induce CSR. Indeed, we found a marked reduction of CSR to IgG1 upon treatment with ASOs masking constitutive I exon dss on GLTs produced in the donor Sμ or the acceptor Sγ1 regions, even though alternative splicing events could be readily detected. By contrast, Kuzin and collaborators showed normal surface Ig expression and serum Ig levels in ΔIμ-s^-/-^ mice harbouring a deletion of 236 bp spanning the constitutive Iμ dss ^28^. As expected, there were no detectable spliced Iμ transcripts in these mice. Nevertheless, alternative “Iμ-like” GLTs driven by regulatory elements other than Eμ were detected at low levels in B cells from ΔIμ-s^-/-^ mice. The authors suggested that such alternative “Iμ-like” GLTs directly contribute to CSR activity ^28^. However, our results showed that these alternative “Iμ-like” GLTs might not be sufficient for efficient CSR activity *in vitro*. Indeed, after treatment of B cells with Iμ dss ASO, we detected Iμ-Cμ alternative transcripts indicative of the use of the same cryptic Sμ dss than described by Kuzin and collaborators whereas a drastic inhibition of CSR was observed. The regulation of μ locus is complex and further investigations are needed to understand the discrepancies concerning the impact of Iμ dss absence at the DNA level, in genetically modified mouse model, and at the RNA level, with our ASO strategy, on CSR. Nonetheless, our data are in agreement with a study of Ruminy and collaborators ^39^. They detected recurrent acquired mutations at the Iμ dss on the functional *IgH* allele of t(14;18) positive lymphomas cases presenting restricted IgM expression and proposed that disruption of Iμ constitutive dss, inducing the expression of abnormal GLTs, may be involved in the perturbation of CSR observed in these lymphomas. Moreover, Spt4 depletion by siRNA in the CH12F3-2A B cell line severely impaired CSR to IgA despite a dramatically increased expression of cryptic transcripts initiating from the Sμ intronic region ^35^. As observed in the donor Sμ region, masking the constitutive I exon dss on GLTs produced in the acceptor Sγ1 region with Iγ1 dss ASO strongly inhibited CSR to IgG1 despite Iγ1-Cγ1 alternative transcripts were detectable. Additional molecular studies will be required to delineate the mechanism involved in the ASO-mediated inhibition of CSR. Several scenarios could explain the inhibition of CSR to IgG1 by I exon dss ASOs. First, RNA-binding proteins have been shown to interact with AID ^22,40–42^ and the ASO, attached to the GLTs, could avoid fixation of such proteins necessary for CSR. Second, it has been described that, when the accumulation of intronic switch RNAs was prevented from the onset of CH12F3-2A B cells stimulation, R-loop levels were decreased by half in S regions ^24^. In our study, after I exon dss ASO treatment, the accumulation of unspliced GLTs indicated a strong splicing inhibition. Thus, even though low levels of constitutive and alternative spliced GLTs are detected after ASO treatment, the lariat abundance may be too weak to induce R-loop formation at S regions and efficient CSR. Third, even if alternatively spliced GLTs were detected after ASOs treatments, the sequence of intronic lariat generated after alternative splicing could prevent efficient CSR. For example, alternative lariat sequence could impede the lariat debranching by DBR1 and the subsequent generation of G-quadruplex RNA structures that participate in guiding AID to specific S regions through RNA:DNA base-pairing ^23^. Alternative lariat sequence could also avoid the fixation of DDX1 to G-quadruplex switch RNAs and consequently impair AID binding to S regions DNA ^24^. These explanations are consistent with a model whereby processed S region transcripts serve as guide RNAs to target AID, in a sequence-dependent manner, to the S regions DNA from which they were transcribed ^36^.

In summary, our study highlighted a dual role for I exon dss during CSR, at the DNA and RNA levels, and further paves the way for antisense strategies in the modulation of antibody class switching and immune response efficiency.

## MATERIALS AND METHODS

### Mice

Two to six month-old *C57BL/6, s-hMT* and *hMT* mice were used. *hMT* and *s-hMT* mice were kindly obtained from Dr. A. Radbruch (Leibniz Institute, Berlin, Germany). Mice were housed and procedures were conducted in agreement with european directive 2010/63/EU on animals used for scientific purposes applied in France as the ‘Décret n°2012-118 du 1er février 2013 relatif à la protection des animaux utilisés à des fins scientifiques’. Accordingly, the present project APAFIS#15279-2018052915087229 v3 was authorized by the ‘Ministère de l’Education Nationale, de l’Enseignement Supérieur et de la Recherche’.

### Splenic B cell *in vitro* stimulation and ASO treatments

Splenic B cells isolated from *C57BL/6, hMT* and *s-hMT* mice were purified with the EasySep Mouse B Cell Isolation Kit (Stemcell Technologies). B cells were cultured for 2 to 4 days in RPMI 1640 with UltraGlutamine (Lonza) containing 10% fetal calf serum (FCS) (Dominique Dutscher), 1 mM sodium pyruvate (Eurobio), 1% AANE (Eurobio), 50 U/ml penicillin / 50 μg/ml streptomycin (Gibco) and 129 μM 2-mercaptoethanol (Sigma-Aldrich). Splenic B cells were stimulated with either 1 μg/ml LPS (LPS-EB Ultrapure, InvivoGen), 1 μg/ml LPS + 20 ng/ml IL4 (Recombinant Murine IL-4, PeproTech) or 5 μg/ml anti-CD40 (mouse CD40/TNFRSF5 MAb (Clone 1C10), Biotechne, USA) + 40 ng/ml IL4. For ASO treatments, vivo-morpholino ASOs (Iγ1 dss ASO: 5’-CCCACTCCCCTGGTCACTTACCG-3’; Iμ dss ASO: 5’-GGCTGCCTCTGGCTTACCATTTG-3’) and an irrelevant ASO (control: 5′-CCTCTTACCTCAGTTACAATTTATA-3′) were designed and purchased at Gene Tools, LLC (Philomath, USA). Stimulated splenic B cells were cultured in the presence of 2 μM ASO. Culture samples were harvested at day 2, 3 or 4 for subsequent flow cytometry analysis, ChIP assays or RNA extraction. Supernatants of stimulated B cells were collected at day 3 or 4 and stored at −20°C until used for ELISA assays.

### Flow cytometry

Cell suspensions of *in vitro* stimulated splenic B cells were washed in phosphate-buffered saline (PBS). To reduce Fc receptor-mediated binding by antibodies of interest, B cells were pre-incubated during 15 minutes with 0.5 μg/ml anti-mouse CD16/CD32 (Clone 2.4G2, BD Pharmingen, ref 553142) in FACS buffer (PBS supplemented with 2% FCS and 2 mM (Ethylenedinitrilo)tetraacetic acid (EDTA) (Sigma-Aldrich)). Cells were then labelled with 0.5 μg/ml anti-mouse B220-BV421 (clone RA3-6B2, BioLegend, ref 103240) and 0.5 μg/ml anti-mouse IgG1-FITC (clone A85-1, BD Pharmingen, ref 553443) antibodies. After 45 minutes, cells were washed in PBS and suspended in FACS buffer. For non-viable cells exclusion, 5 μl of 7-AAD (BD Pharmingen, ref 559925) were added on cells 10 minutes before flow cytometry analysis. Data were acquired on a BD Pharmingen Fortessa LSR2 (BD Biosciences, San Jose, CA, USA) and analysed using Flowlogic™ software (Miltenyi Biotec).

### ELISA assays

Culture supernatants and sera were analysed for the presence of IgM, IgG1 or IgG2b by ELISA. Blood samples were collected from 12 weeks-old *hMT* and *s-hMT* mice. Serum samples were recovered by centrifugation and stored at −20°C until used. ELISA assays were performed in polycarbonate 96 multiwell plates, coated overnight at 4°C (100 μl per well) with 2 μg/ml IgM, IgG1 or IgG2b antibodies (Southern Biotechnologies: goat anti-Mouse IgM human ads UNLB, ref 1020-01; goat anti-Mouse IgG1 Human ads-UNLB, ref 1070-01; goat anti-Mouse IgG2b Human ads-UNLB, ref 1090-01) in PBS. After three successive washing steps in PBS with 0.05% Tween® 20 (Sigma-Aldrich), a blocking step with 100 μl of 3% bovine serum albumin (BSA) (Euromedex) in PBS was performed for 30 min at 37°C. After three washing steps, 50 μl of sera / supernatant, or standards IgM, IgG1 or IgG2b (Southern Biotechnologies, 400 ng/ml) were diluted into successive wells in 1% BSA/PBS buffer and incubated for 2 h at 37°C. After three washing steps, 100 μl per well of 1 μg/ml Alkaline Phosphatase (AP)-conjugated goat anti-mouse antibodies (Southern Biotechnologies: goat anti-Mouse IgG1 Human ads-AP, ref 1070-04 and goat anti-mouse kappa-AP, ref 1050-04; Cell Lab: goat anti-Mouse IgG2b Human ads-AP, ref 731943) were incubated in PBS with 0.05% Tween® 20 for 2 h at 37°C. After three washing steps, AP activity was assayed: 100 μl of substrate for AP (SIGMA*FAST* ™ p-Nitrophenyl phosphate Tablets, Sigma-Aldrich) were added and, after 15 min, the reaction was blocked with addition of 50 μl of 3 M NaOH (Sigma-Aldrich). Optic density was then measured at 405 nm on a Multiskan FC microplate photometer (Thermo Scientific).

### RT-PCR and quantitative RT-PCR

*In vitro* stimulated splenic B cells were harvested and RNA was extracted using TRIzol™ Reagent (Invitrogen) procedure. Reverse transcription was carried out on 1 μg of DNase I (Invitrogen)-treated RNA using High-Capacity cDNA Reverse Transcription kit (Applied Biosystems). Priming for reverse transcription was done with random hexamers.

To analyse GL transcription, PCRs were performed on cDNA using the Taq Core Kit (MP Biomedicals) and appropriate primer pairs (primer pairs are provided in Supplementary table 1). PCR amplification of the -actin transcript was used as an internal loading control.

To determine nucleotide sequence of normal and alternative transcripts, PCRs were performed on cDNA using the Phusion^®^ High-Fidelity DNA Polymerase (New England BioLabs) and appropriate primer pairs. After purification of the RT-PCR products using the NucleoSpin Gel and PCR Clean-up kit (Macherey-Nagel) according to the manufacturer’s instructions, sequencing was performed using the BigDye™ Terminator v3.1 Cycle Sequencing Kit on a 3130xl Genetic Analyzer ABI PRISM (Applied Biosystems). Quantification of the purified RT-PCR products was also performed using an Agilent 2100 Bioanalyzer (Agilent Technologies) according to the Agilent High Sensitivity DNA kit instructions.

Quantitative PCRs were performed on cDNA using Premix Ex Taq™ (probe qPCR), ROX Plus (Takara) or TB Green Premix Ex Taq™ II (Tli RNase H Plus), ROX Plus (Takara) on a StepOnePlus Real-Time PCR system (Applied Biosystems). Transcripts were quantified according to the standard 2^−ΔΔCt^ method after normalization to *Gapdh*. Primers and probes used for determination of transcripts are listed in Supplementary table 1.

### ChIP assays

ChIP assays were performed using anti-H3ac (Millipore, 06-599), anti-H3K4me3 (Millipore, 07-473), anti-H3K9ac (Millipore, 06-942), anti-RNA pol II (CTD4H8, Santa Cruz Biotechnology, sc-47701), anti-RNA pol II ser2P (Abcam, ab5095), and anti-RNA pol II ser5P (Abcam, ab5131) as previously described ^43^. In brief, 1×10^7^ LPS stimulated B cells from *hMT* and *s-hMT* mice were harvested at day 2, washed twice in PBS and cross-linked at 37°C for 15 min in 15 ml of PBS with 1% formaldehyde (Sigma-Aldrich). The reaction was quenched with 0.125 M glycine (Sigma-Aldrich). After lysis, chromatin was sonicated to 0.5–1 kb using a Vibracell 75043 (Thermo Fisher Scientific). After dilution in ChIP buffer (0.01% SDS (Sigma-Aldrich), 1.1% Triton X-100 (Sigma-Aldrich), 1.2 mM EDTA (Eurobio), 16.7mM Tris-HCl (Sigma-Aldrich), pH 8.1, and 167 mM NaCl (Sigma-Aldrich)), chromatin was precleared by rotating for 2 h at 4°C with 100 μl of 50% protein A/G slurry (0.2 mg/ml sheared salmon sperm DNA, 0.5 mg/ml BSA, and 50% protein A/G; Sigma-Aldrich), 0.3−0.5×10^6^ cell equivalents were saved as input, and 3−5×10^6^ cell equivalents were incubated overnight with specific or control antibodies. Immune complexes were precipitated by the addition of protein A/G. Cross-linking was reversed by overnight incubation (70°C) in Tris-EDTA (Sigma-Aldrich) buffer with 0.02% SDS, and genomic DNA was obtained after phenol/chloroform extraction. Analysis of immuno-precipitated DNA sequences was done by quantitative PCR using the primer pairs described in Supplementary table 1.

### Statistical analysis

Results are expressed as means ± SEM and overall differences between variables were evaluated by an unpaired two-tailed Student’s *t* test using Prism GraphPad software (San Diego, CA).

## ACKNOWLEDGEMENTS

We thank the staff of our animal facility. We also thank J. Cook-Moreau and E. Pinaud (UMR CNRS 7276 – INSERM 1262, Limoges, France) for critical reading of the manuscript and A. Radbruch (Leibniz Institute, Berlin, Germany) for providing *hMT* and *s-hMT* mice. This work was supported by grants from Fondation ARC (PJA 20161204724), INCa (PLBIO15-256), ANR (2017-CE15-0024-01), Ligue Contre le Cancer (CD87, CD19, CD23), and Fondation Française pour la Recherche contre le Myélome et les Gammapathies monoclonales (FFRMG).

## AUTHOR CONTRIBUTIONS

AM and MOA performed experiments and analysed data. JML and NS performed experiments. SLP and LD conceived the project, designed experiments, analysed data and wrote the manuscript.

## SUPPLEMENTARY MATERIAL

The online version of this article contains supplementary material.

## FIGURE LEGENDS

**Supplementary figure 1. Defect of IgG1 class switching in mice lacking Iγ1 dss**

(A) Quantification of Ig isotypes (IgM, IgG2b, and IgG1) in sera of homozygous *s-hMT* and *hMT* mice by ELISA. (B-D) Splenic B cells were isolated from homozygous *s-hMT* and *hMT* mice and stimulated with LPS. After 4 days stimulation, amounts of Ig isotypes (IgM, IgG2b, and IgG1) were determined in culture supernatants by ELISA (B). After 3 days stimulation, post-switch Iμ-Cγ1 (C) and AID (D) mRNA expression relative to GAPDH mRNA expression was monitored by quantitative RT-PCR. Expression of Iμ-Cγ1 or AID in B cells from *s-hMT* mice was normalized to 1. Data are means ± SEM, n=3 to 4 for each genotype. Unpaired two-tailed Student’s t test was used to determine significance. ND: not detected, ns: non significant, **** P < 0.0001.

**Supplementary figure 2. Similar RNA pol II binding in Sμ and Sγ2b regions of *s-hMT* and *hMT* mice**

Splenic B cells were isolated from homozygous *s-hMT* and *hMT* mice and stimulated with LPS. After 2 days, the cells were analysed for Ser2P RNA pol II (A, B) and Ser5P RNA pol II (C, D) levels in Sμ (A, C) and Sγ2b (B, D) regions by ChIP coupled to quantitative PCR. Background signals from mock samples with irrelevant antibody were subtracted. Values were normalized to total input DNA. Primers (triangles) used for quantitative PCR are described on the illustrative schema (bottom). Data are means ± SEM of at least two independent experiments, n=4 for each genotype. Unpaired two tailed Student’s t test was used to determine significance. ns: non significant.

**Supplementary figure 3. Sequences of γ1 constitutive and alternative spliced transcripts**

The sequences of Iγ1 exon (bold) and CH1γ1 exon are indicated. Donor (red) and acceptor (green) splice sites are also represented.

**Supplementary table 1. Primers used for ChIP, RT-PCR and quantitative RT-PCR experiments**

