## supplementary figures and table for "Uncoupling splicing from transcription using antisense oligonucleotides reveals a dual role for I exon donor splice sites in antibody class switching"

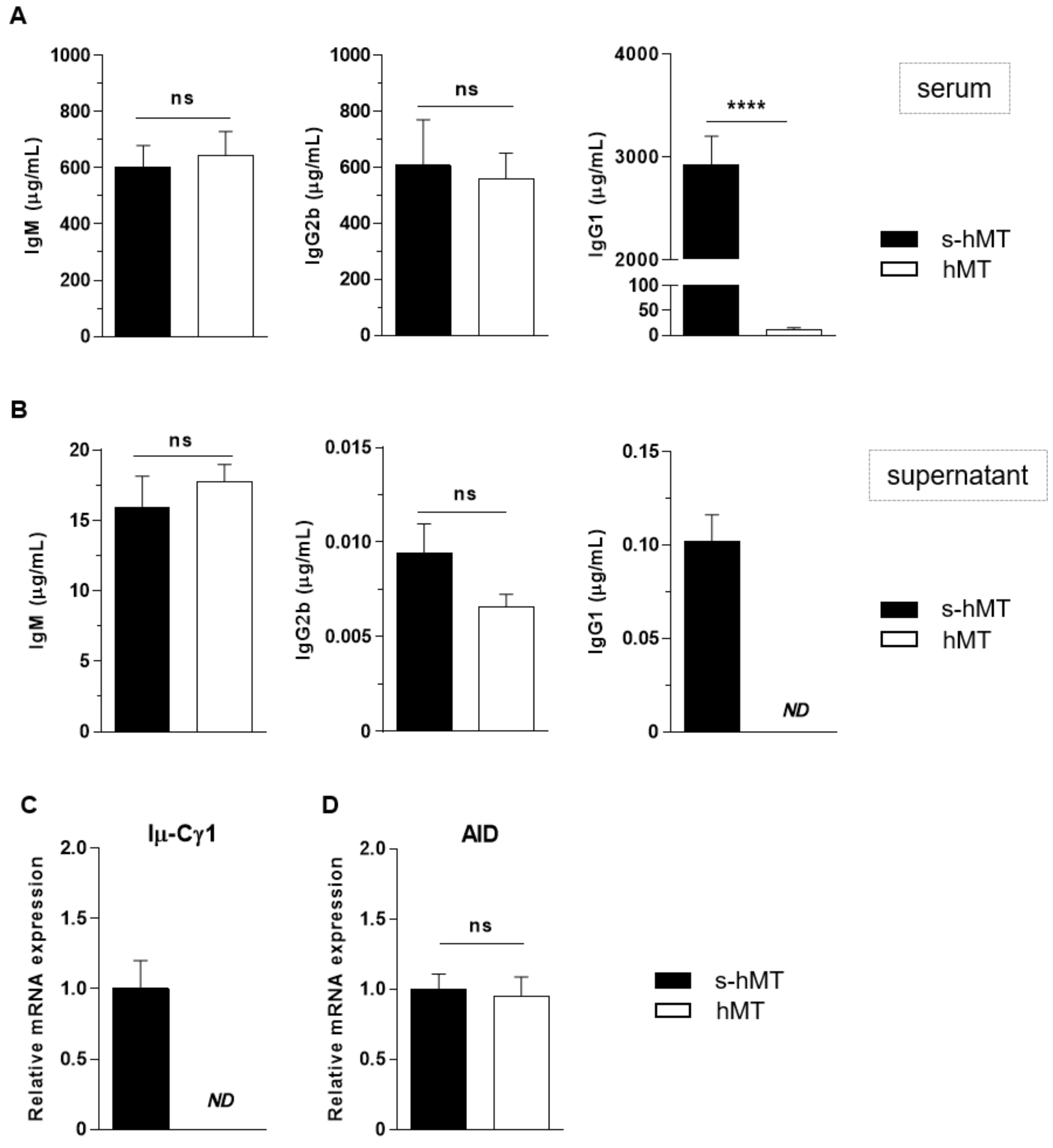


**Supplementary figure 1. Defect of IgG1 class switching in mice lacking Iγ1 dss**

(A) Quantification of Ig isotypes (IgM, IgG2b, and IgG1) in sera of homozygous *s-hMT* and *hMT* mice by ELISA. (B-D) Splenic B cells were isolated from homozygous *s-hMT* and *hMT* mice and stimulated with LPS. After 4 days stimulation, amounts of Ig isotypes (IgM, IgG2b, and IgG1) were determined in culture supernatants by ELISA (B). After 3 days stimulation, post-switch Iμ-Cγ1 (C) and AID (D) mRNA expression relative to GAPDH mRNA expression was monitored by quantitative RT-PCR. Expression of Iμ-Cγ1 or AID in B cells from *s-hMT* mice was normalized to 1. Data are means ± SEM, n=3 to 4 for each genotype. Unpaired two-tailed Student's t test was used to determine significance. ND: not detected, ns: non significant, **** $P < 0.0001$.

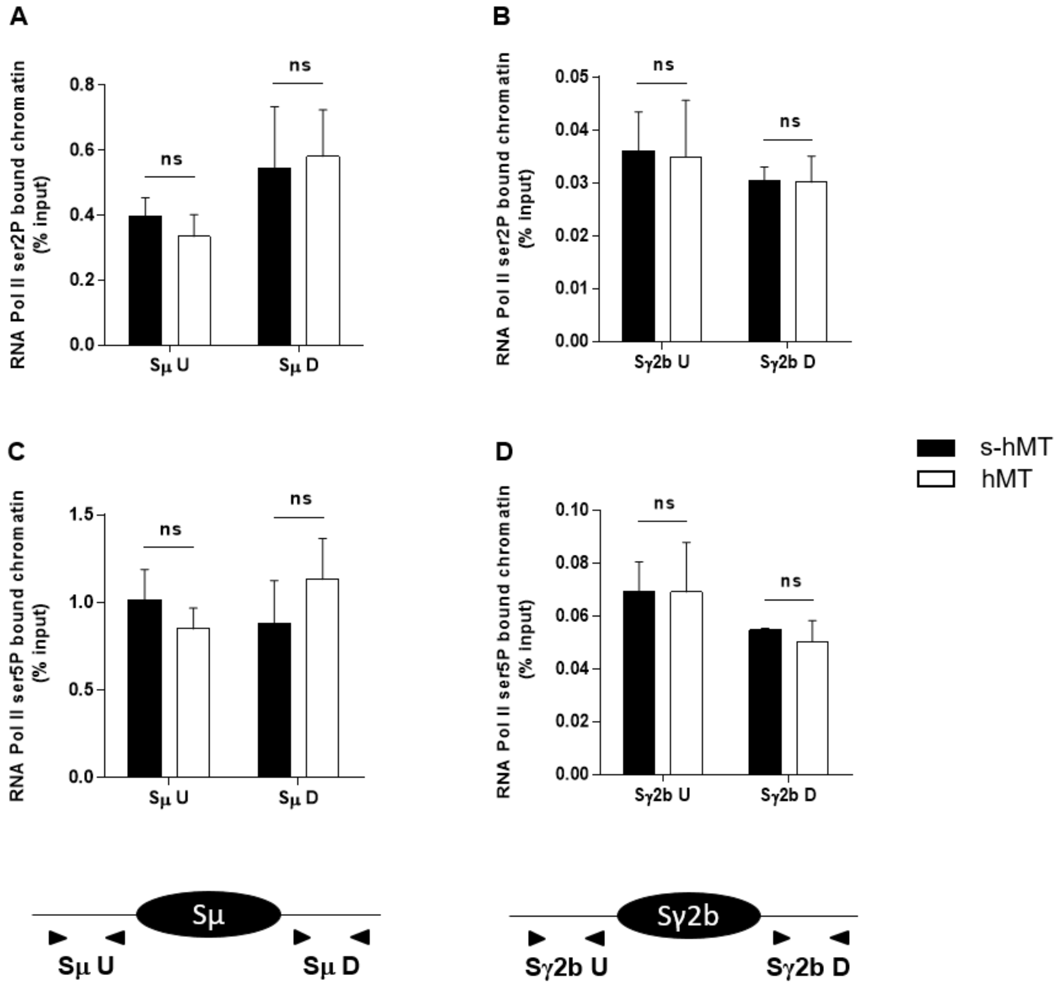


**Supplementary figure 2. Similar RNA pol II binding in Sμ and Sγ2b regions of *s-hMT* and *hMT* mice**

Splenic B cells were isolated from homozygous *s-hMT* and *hMT* mice and stimulated with LPS. After 2 days, the cells were analysed for Ser2P RNA pol II (A, B) and Ser5P RNA pol II (C, D) levels in Sμ (A, C) and Sγ2b (B, D) regions by ChIP coupled to quantitative PCR. Background signals from mock samples with irrelevant antibody were subtracted. Values were normalized to total input DNA. Primers (triangles) used for quantitative PCR are described on the illustrative schema (bottom). Data are means ± SEM of at least two independent experiments, n=4 for each genotype. Unpaired two tailed Student's t test was used to determine significance. ns: non significant.

**Supplementary figure 3: Sequences of γ1 constitutive and alternative spliced transcripts**

The sequences of Iγ1 exon (bold) and CH1γ1 exon are indicated. Donor (red) and acceptor (green) splice sites are also represented.

**Constitutive γ1 transcript:**

**GTCAATCATATGATGGAAAGAGGGTAGCATTCACCTCTCTGGGACAAAGGCT**
**GTGACTCTGGGAAAGACAAGAGAAGGGCAGGACCAAAACAGGAACAGAGAC**
**GGCTGCTTTCACAGCTTCCACATGTGAGTGGGGTCAGCAGGGAAAGGAGCT**
**GCAAGAAGAGGCCATACAAACAGCACGCATCTGTGGCCCTTCCAGATCTTTG**
**AGTCATCCTATCACGGGAGATTGGGAAGGAGTTGACAGACCAGCCCAGGCA**
**GAGGAAGCCTCTGTGTTAAAGAGTAAAGGTGCTTGCCTACAGCCTGGTGTCA**
**ACTAGGCAGGCCCTGGGGGGCCGGGAAGGGGCCTCCTAGACAAGCACAGGC**
**ATGTAGAGCTGCACACCCCACAGACAAACCTGAGCCCCGAGGATATCATGG**
**AATATATCGAGAAGCCTGAGGAATGTGTTTGGCATGGACTACAGGTTGAGAG**
**AACCAAGGAAGCTGAGCCCTGCG**CCAAAACGACACCCCCATCTGTCTATCCAC
TGGCCCCTGGATCTGCTGCCCAAACTAACTCCATGGTGACCCTGGGATGCCTGGT
CAAGGGCTATTTCCCTGAGCCAGTGACAGTGACCTGGAACTCTGGATCCCTGTCC
AGCGGTGTGCACACCTTCCCAGCTGTCCTGCAGTCTGACCTCTACACTCTGAGCA
GCTCAGTGACTGTCCCCTCCAGCACCTGGCCCAGCCAGACCGTCACCTGCAACGT
TGCCCACCCGGCCAGCAGCACCAAGGTGGACAAGAAAATTG

**Alternative γ1 transcript 1:**

**GTCAATCATATGATGGAAAGAGGGTAGCATTCACCTCTCTGGGACAAAGGCT**
**GTGACTCTGGGAAAGACAAGAGAAGGGCAGGACCAAAACAGGAACAGAGAC**
**GGCTGCTTTCACAGCTTCCACATACAAACCTGAGCCCCGAGGATATCATGGA**
**ATATATCGAGAAGCCTGAGGAATGTGTTTGGCATGGACTACAGGTTGAGAGA**
**ACCAAGGAAGCTGAGCCCTGCG**CCAAAACGACACCCCCATCTGTCTATCCACT
GGCCCCTGGATCTGCTGCCCAAACTAACTCCATGGTGACCCTGGGATGCCTGGTC
AAGGGCTATTTCCCTGAGCCAGTGACAGTGACCTGGAACTCTGGATCCCTGTCCA
GCGGTGTGCACACCTTCCCAGCTGTCCTGCAGTCTGACCTCTACACTCTGAGCAG
CTCAGTGACTGTCCCCTCCAGCACCTGGCCCAGCCAGACCGTCACCTGCAACGTT
GCCCACCCGGCCAGCAGCACCAAGGTGGACAAGAAAATTG

**Alternative γ1 transcript 2:**

**GTCAATCATATGATGGAAAGAGGGTAGCATTCACCTCTCTGGGACAAAGGCT**
**GTGACTCTGGGAAAGACAAGAGAAGGGCAGGACCAAAACAGGAACAGAGAC**
**GGCTGCTTTCACAGCTTCCACAT**CCAAAACGACACCCCCATCTGTCTATCCACT
GGCCCCTGGATCTGCTGCCCAAACTAACTCCATGGTGACCCTGGGATGCCTGGTC
AAGGGCTATTTCCCTGAGCCAGTGACAGTGACCTGGAACTCTGGATCCCTGTCCA
GCGGTGTGCACACCTTCCCAGCTGTCCTGCAGTCTGACCTCTACACTCTGAGCAG

CTCAGTGACTGTCCCCTCCAGCACCTGGCCCAGCCAGACCGTCACCTGCAACGTT
GCCCACCCGGCCAGCAGCACCAAGGTGGACAAGAAAATTG

**Alternative γ1 transcript 3:**

**GTCAATCATATGATGGAAAGAGGGTAGCATTCACCTCTCTGGGACAAAGGCT
GTGACTCTGGGAAAGACAAGAGAAGGGCAGGACCAAAACAGGAACAGAGAC
GGCTGCTTTCACAGCTTCCACATGTGAGTGGGGTCAGCAGGGAAAGGAGCT
GCAAGAAGAGGCCATACAAACAGCACGCATCTGTGGCCCTTCCAGATCTTTG
AGTCATCCTATCACGGGAGATTGGGAAGGAGTTGACAGACCAGCCCAGGCA
GAGGAAGCCTCTGTGTTAAAGAGTAAAG**CCAAAACGACACCCCCATCTGTCTA
TCCACTGGCCCCTGGATCTGCTGCCCAAACTAACTCCATGGTGACCCTGGGATGC
CTGGTCAAGGGCTATTTCCCTGAGCCAGTGACAGTGACCTGGAACTCTGGATCCC
TGTCCAGCGGTGTGCACACCTTCCCAGCTGTCCTGCAGTCTGACCTCTACACTCTG
AGCAGCTCAGTGACTGTCCCCTCCAGCACCTGGCCCAGCCAGACCGTCACCTGCA
ACGTTGCCCACCCGGCCAGCAGCACCAAGGTGGACAAGAAAATTG

### Supplementary table 1: Primers used for ChIP, RT-PCR and quantitative RT-PCR experiments

| Name | Sequence | ChIP | RT-PCR | qRT-PCR |
|---|---|---|---|---|
| hMT promoter-for | 5' CCCGGTCTCTCGAGCTATAAAC 3' | x | | |
| hMT promoter-rev | 5' GGTTCGCTGGGACTTGGA 3' | x | | |
| hMT promoter-Probe | 5' CTGCTTGCATGTGGAATTGTGAGCG 3' | x | | |
| Sγ1-U-for | 5' AGGACACAAGACCTGCAAAAGAG 3' | x | | x |
| Sγ1-U-rev | 5' CCCAGGAGCTGCTGAACCT 3' | x | | x |
| Sγ1-U-Probe | 5' TGAGGCTGGTAAGAGTAACAAGGTAACCTGGG 3' | x | | x |
| Sγ1-D-for | 5' CAGGCAAACTAAACCAGTGGG 3' | x | | |
| Sγ1-D-rev | 5' AGGATGTCCACCCTCACCCAGGC 3' | x | | |
| Sμ-U-for | 5' TCTAAAATGCGCTAAACTGAGG 3' | x | | |
| Sμ-U-rev | 5' AGCGTAGCATAGCTGAGCTC 3' | x | | |
| Sμ-D-for | 5' CTGAATGAGTTTCACCAGGCC 3' | x | | |
| Sμ-D-rev | 5' GGCCTGTCCTGCTTGGCTTC 3' | x | | |
| Sγ2b-U-for | 5' AGCTCCAAAAGCTCAGCAGAC 3' | x | | |
| Sγ2b-U-rev | 5' AGCCCCAGCTTACAAAGAGCT 3' | x | | |
| Sγ2b-D-for | 5' GGTGGGAATATGAGGGAGAAGTCCTAG 3' | x | | |
| Sγ2b-D-rev | 5' TTCCACCTGCCTCAGCTCTCCCACAGC 3' | x | | |
| Iμ-for | 5'-TTGACATTCTGGTCAAAACGGC | | x | |
| Cμ-rev | 5'-TCTGAACCTTCAAGGATGCTCTTG | | x | |
| Iγ1-for | 5'-CCAAAACAGGAACAGAGACGG | | x | |
| Cγ1-rev | 5'-TAGACAGATGGGGGTGTCGT | | x | |
| Actin-for | 5' CGATGCCCTGAGGCTCTT 3' | | x | |
| Actin-rev | 5'-TAAAACGCAGCTCAGTAACAGTCCG | | x | |
| Iμ-for-Q | 5'-ACCTGGGAATGTATGGTTGTGGCTT | | | x |
| Iγ1-for-Q | 5'-GAACCAAGGAAGCTGAGCCC | | | x |
| Iε-for-Q | 5'-AGATTCACAACGCCTGGGAG | | | x |
| Cγ1-rev-Q | 5'-ATGGAGTTAGTTTGGGCAGCA | | | x |
| Cε-rev-Q | 5'-AATACCAGGTCACAGTCACAGG | | | x |
| Sγ1U-rev-Q | 5'-AATGCTGGGATTGATCCTGGG | | | x |
| SεU-rev-Q | 5'-GCAAACCCTTTTGCTCAGGG | | | x |
| Gapdh-probe | Mm99999915_g1 (Life Technologies) | | | x |
| AID-probe | Mm01184115_m1 (Life Technologies) | | | x |
